# Urbanization alters ecological and evolutionary interactions between Darwin’s finches and *Tribulus cistoides* on the Galápagos Islands

**DOI:** 10.1101/2020.09.26.301507

**Authors:** L. Ruth Rivkin, Reagan A. Johnson, Jaime A. Chaves, Marc T.J. Johnson

## Abstract

Emerging evidence suggests that urbanization shapes the ecology and evolution of species interactions. Islands are particularly susceptible to urbanization due to the fragility of their ecosystems; however, few studies have examined the effects of urbanization on species interactions on islands. To address this gap, we studied the effects of urbanization on interactions between Darwin’s finches and its key food resource, *Tribulus cistoides*, in three towns on the Galápagos Islands. We assessed the effects of urbanization on seed and mericarp removal, mericarp morphology, and finch community composition using natural population surveys, experimental manipulations, and finch observations. We found that both seed and fruit removal rates were higher in urban compared to non-urban populations in the natural and experimental populations, and that urbanization modified selection on mericarp size and defense. Urban environments supported smaller and less diverse finch communities than non-urban environments. Together, our results suggest that urbanization can dramatically alter ecological interactions between Darwin’s finches and *T. cistoides*, leading to modified selection on *T. cistoides* populations. Our study demonstrates that urban development on islands can have profound effects on the ecology and evolution of trophic interactions.

## 1. Introduction

Urbanization results in substantial changes to the environment. Urban habitats are typically warmer, more polluted, and more fragmented than nearby non-urban habitats, which can lead to changes in the abundance and persistence of populations, and altered diversity and community [1–3]. Emerging evidence suggests that ecological changes associated with urbanization may alter natural selection and drive the evolution of novel adaptations [4,5]. Most examples of contemporary urban evolution occur in well-established cities, especially in Europe and North America [4,6]. However, we still have limited knowledge of how urbanization in tropical regions and particularly on islands can influence the ecology and evolution of species. Islands may be particularly sensitive to urbanization because of the unique and often fragile ecosystems they support, and even small human settlements may have large-scale effects on the ecosystem [7]. Our study addresses these gaps using the iconic Darwin’s finch-*Tribulus* interaction of the Galápagos archipelago.

Urbanization can affect both the ecology and evolution of species interactions [8]. Predictions about how interactions will respond to urbanization are complex, especially for antagonistic interactions [9,10]. Antagonistic (e.g. predator-prey) interactions are inherently interconnected, and may be susceptible to urbanization through effects on one or both trophic levels. For instance, urbanization may decouple predator-prey interactions through the addition of food subsidies from anthropogenic resources [11], or may be intensify interactions when urban habitat fragmentation reduces available niche space, increasing the frequency of interactions [12,13]. These changes may lead to novel selection pressures on one or both interacting species [10]. Despite increasing work on species interactions in urban environments, it remains unclear how urbanization simultaneously shapes both the ecology and evolution of these interactions.

The Galápagos Islands of Ecuador provide an ideal system to test questions about how urbanization affects the ecology and evolution of species interactions on islands. Ground finches and *Tribulus cistoides* L. (Zygophyllaceae, common names puncture vine or Jamaican feverplant)) on the Galápagos have a long history of study [19–21], and it is clear that these species are experiencing a dynamic and ongoing co-evolutionary arms-race [17]. When resources are scarce during the dry season, *T. cistoides* is an important food resource for three medium and large beaked ground finch species: *Geospiza fortis, G. magnirostris*, and *G. conirostris* [18–20]. Predation on *T. cistoides* has led niche segregation and evolutionary changes in beak morphology in these finch species [21–23]. In turn, finches influence mericarp survival, and select for smaller, harder, and more defended mericarps [17]. Urbanization on the Galápagos Islands influences both finches and *T. cistoides* populations [24–28]. Humans are one of the primary dispersers of *T. cistoides* on the Galápagos [27], and urbanization reduces resource partitioning through selection on beak morphology in *G. fortis* [25,29], likely due to the increased availability of human food [25] and urban-induced behaviour modifications [30].

Despite clear evidence of the impacts of urbanization on ground finches and *T. cistoides* individually, no work has examined the effects of urbanization on interactions between these species. Our objective was to understand the effects of urbanization on the ecology and evolution of interactions between ground finches and *T. cistoides*. We studied seed removal rates, selection on mericarp morphology, and ground finch community composition on three islands on the Galápagos archipelagos (figure 1). We used this system to ask three questions: 1) Does seed removal by finches differ between urban and non-urban populations of *T. cistoides*? 2) Does urbanization alter selection imposed by seed removal on mericarp size and defense? 3) Does the abundance and community structure of Galápagos finch populations differ between urban and non-urban sites, and how might this relate to patterns of seed removal and selection? Here, we aim to identify the joint effects of how urbanization affects the ecology and evolution of antagonistic species interactions in a fragile island ecosystem.

**Figure 1.**
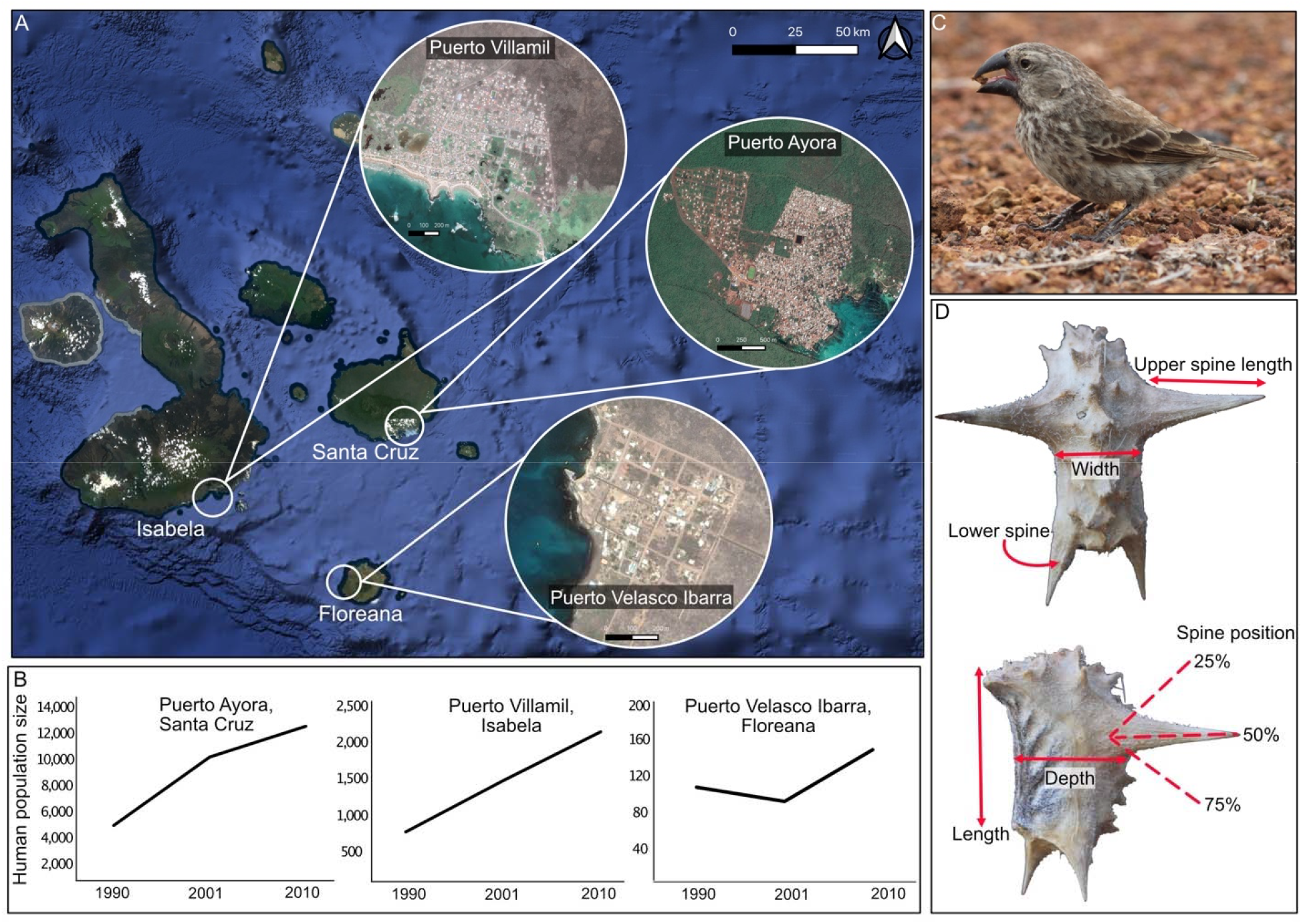
a) Map of the Galápagos Islands, with the three islands sampled and their principle towns. Maps were taken from Google Satellite dating from 2018. B) Change in population size in each town from 1990-2010 [41], ordered from largest (Santa Cruz) to smallest (Floreana); note, the human population has continued to grow rapidly but censuses are carried out only every 10 years. c) A female medium ground finch (*Geospiza fortis*) holding a *Tribulus cistoides* mericarp in its beak. d) Dorsal and lateral images of a *T. cistoides* mericarp, with each of the six morphological traits measured. All images taken by MTJJ.

## 2. Methods

### a) Study site and system

We studied seed and fruit removal rates, selection on mericarp morphology, and finch community composition on three islands on the Galápagos archipelagos: Floreana, Isabela, and Santa Cruz. We selected these locations because both large beaked ground finches and *T. cistoides* are common in and around the towns. We have described our study site and study system in detail in the supplemental methods.

### b) Study design

This study was comprised of three components: a survey of natural populations of *T. cistoides* to estimate seed removal rates; an experiment with mericarp defense traits artificially manipulated to measure mericarp removal; and ground finch community observations. We studied each of these components at the end of the dry season from January-March 2018 in urban and non-urban locations on each of the three islands. For each component described below, we considered a population to be urban if it occurred within a town’s borders.

#### Natural population survey

To test for differences in seed removal rates and selection among urban and non-urban populations, we conducted a survey of seed removal in natural *T. cistoides* populations in February 2018. Natural populations provided us with a picture of natural variation in seed removal between urban and non-urban habitats across islands over 8-12 months because mericarps are produced in the wet season, and typically persist for many months until germinating or decaying the following wet season [31]. We sampled 16 populations on Floreana (*N* = 9 urban and 7 non-urban), 28 populations on Isabela (*N* = 15 urban and 13 non-urban), and 41 populations on Santa Cruz (*N* = 22 urban and 19 non-urban). We collected 20 mericarps from each population (except for one population where we only found 19 mericarps), for a total of 1,699 mericarps. We counted the number of seeds eaten from each mericarp to estimate seed removal rate. Following a modified protocol outlined in Carvajal et al. [17], we measured six size and defense traits on each mericarp: mericarp length, width, depth, the length of the longest spine, presence or absence of lower spines, and spine position (figure 1d).

#### Fruit removal experiment

At the same time as the natural population surveys, we conducted a six week-long experiment to test for variation in fruit removal rates and selection in *T. cistoides*. This experiment complemented our natural population surveys by allowing us to causally determine how morphology affects removal and natural selection on mericarps by finches. However, the shorter-term nature of the experiment meant that mericarp removal rates provided a shorter window of predation pressure. We collected 800-900 intact mericarps (i.e. not attacked by finches) from non-urban populations on each island. We weighed the mericarps and measured the same six morphological traits measured in the natural population survey (figure 1d). We selected 20 urban and 20 non-urban populations per island (*N* = 40 populations per island) and placed a petri dish (100 mm diameter) in each population. Each dish contained 20 mericarps placed on top of locally collected substrate (i.e. volcanic sand and gravel) for a total of 2,120 mericarps. We randomly selected half of the mericarps and used wire cutters to clip off all of their spines to create an “undefended” mericarp, while the other half were left with their spines intact as “defended” mericarps. Although *T. cistoides* exhibits natural variation in spine number [17,18], we selected mericarps that had four spines so that our manipulation simulated mericarps with four (defended) versus zero (undefended) spines. We marked each mericarp with a unique identifier on its dorsal surface using a black sharpie marker so that we could identify each individual mericarp at the conclusion of the experiment.

We left the mericarps in the field for six weeks, and then collected them to score removal. Using the identifying marks placed on the mericarps, we determined which mericarps had been removed and which remained in the tray. If a mericarp was removed, we counted it as “eaten” because ground finches often carry mericarps away from the location where they collect them to crack them on a hard surface. We also counted the number of seeds removed from each mericarp that was recovered, but the number of recovered mericarps with seeds was too small for analysis (< 1% of the total sample), so our analyses focused on the removal rate. We carefully placed petri dishes in locations where humans would not walk, thus we are confident that mericarp removal was due to finch consumption and not human dispersal [27]. Some petri dishes were disturbed during the experiment (three on Floreana, six on Isabela, eight on Santa Cruz), and these petri dishes were excluded from the analyses.

#### Finch community observations

To determine how ground finch community composition varies with urban development, we conducted surveys at urban and non-urban *T. cistoides* natural populations on each island. We selected six sites per island (*N* = 3 urban and 3 non-urban) ensuring that each site had clear lines of sight within 50 m of the center of the population. We surveyed each location for five minutes and recorded finch sightings within 50 m during that time. Although *G. fortis* and *G. magnirostris* are the only vertebrate seed predators of *T. cistoides* seeds on the islands we studied [17], we recorded all finch species that frequently interact with *G. fortis* and *G. magnirostris* and thus could influence their distribution, abundance, or behavior. We repeated the surveys four times on Santa Cruz and three times on Floreana and Isabela.

### c) Statistical analyses

We used R v 3.6.2 [32] for all analyses, details about the statistical analyses can be found in the supplemental materials and methods. The R code and data files can be found in the supplemental materials.

## 3. Results

### a) Natural population survey

There were effects of urbanization, mericarp size, and defense on seed removal rate in *T. cistoides* populations (Table S2). The number of seeds eaten per mericarp was 1.25% higher in urban populations than in non-urban (*Urbanization*: χ^2^_1_ = 3.91, p = 0.048). More seeds were eaten from small mericarps (*Size*: χ^2^_1_ = 10.74, p < 0.001), but on average seed removal was greater in urban populations than from non-urban population (*Urbanization* × *Size*: χ^2^_1_ = 4.51, p = 0.034; figure 2a), suggesting seed removal imposed stronger selection for large mericarps in urban populations. There was no main effects of island or mericarp defense, however the effect of both size and defense differed among islands *(Island* × *Size*: χ^2^_2_ = 8.89, p = 0.012; *Island* × *Defense*: χ^2^_2_ = 6.69, p = 0.035), consistent with selection on these traits varying among islands.

**Figure 2.**
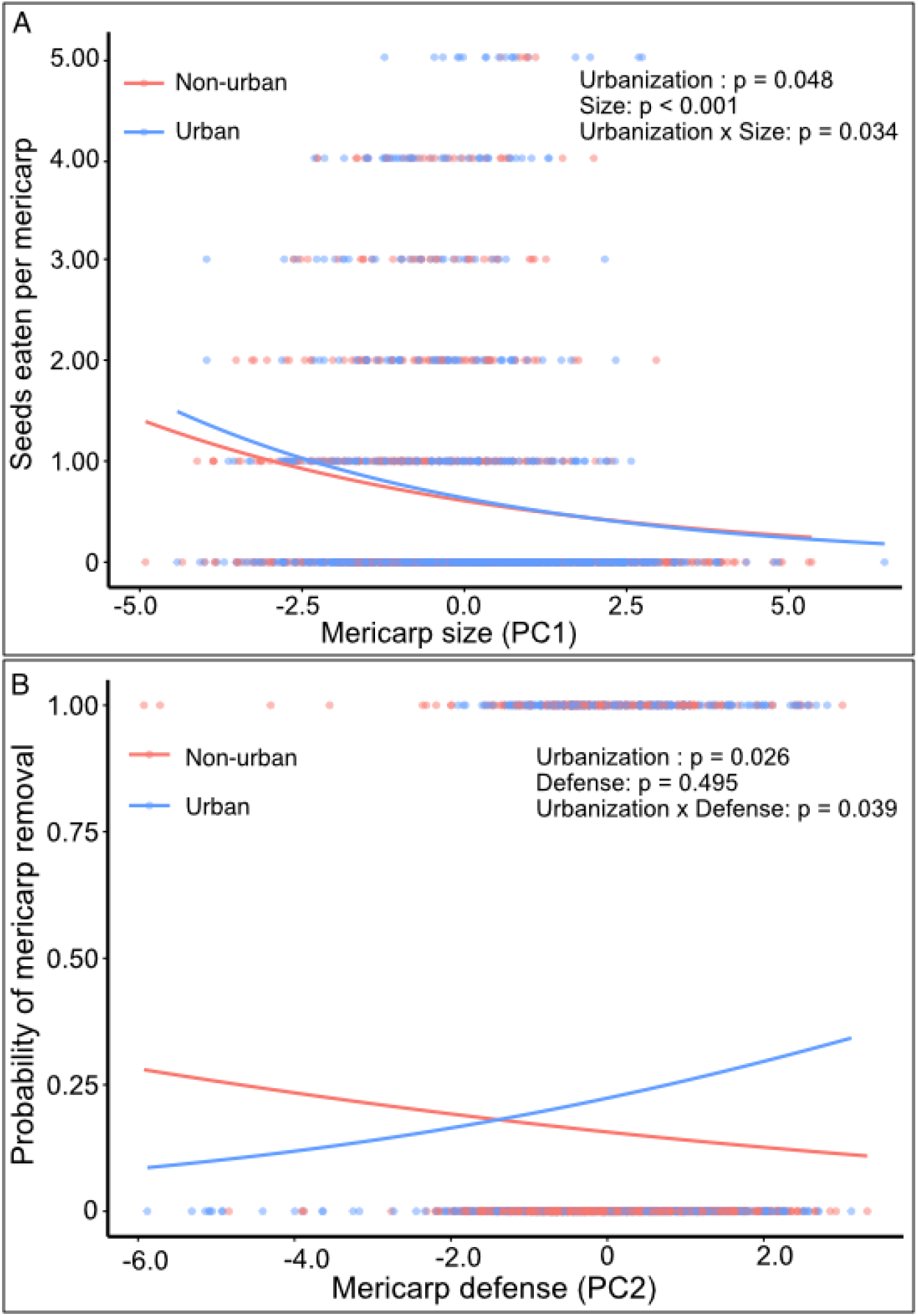
Seed and fruit removal by finches in natural and experimental populations of *Tribulus cistoides*. a) The number of seeds eaten per mericarp declined with cumulative mericarp size (PC1) in natural populations of *T. cistoides*, with small mericarps being eaten more in urban areas than in non-urban areas. b) The probability of mericarp removal from experimental populations increased with mericarp defense (PC2) in urban populations but declined in non-urban populations.

### b) Fruit removal experiment

Mericarp removal during our experiment was influenced by urbanization and defense traits (Table S2). Mericarp removal rate was 43% higher in urban populations (*Urbanization*: χ^2^_1_ = 4.98, p = 0.026), and 39% higher on clipped (undefended) mericarps (*Clipped*: χ^2^_1_ = 8.44, p = 0.004), although this effect did not vary with urbanization (*Urbanization* × *Clipped*: χ^2^_1_ = 0.05, p = 0.823).

However, urbanization interacted with natural variation in mericarp defenses (*Urbanization* × *Defense*: χ^2^_1_ = 4.24, p = 0.039). Well-defended mericarps were removed more often from urban populations compared to non-urban populations (figure 2b), suggesting that fruit removal by finches imposes selection for poorly defended mericarps in urban populations. In contrast, mericarps that were poorly defended experienced a higher removal rate in non-urban populations than in urban populations, indicating that the defensive function of spines switched between urban and non-urban habitats (figure 2b). To better understand which traits were causing this effect, we ran additional analyses that included individual defense traits as covariates, instead of the composite defense trait (supplemental methods). We found that mericarps with longer spines were more likely to be removed in urban populations than non-urban populations (*Urbanization* × *Spine length*: χ^2^_1_ = 4.95, p = 0.027; Table S2), but no effect of lower spine or spine position.

### c) Finch community composition

We observed five species of ground finches across sites: two *G. magnirostris*, 171 *G. fortis*, 268 *G. fuliginosa*, 54 *G. scandens*, and 11 *Platyspiza crassirostris*. We found no change in the total or relative abundances of *G. fortis* in the finch communities among urban and non-urban sites (*Urbanization*: χ^2^_1_ = 1.10, p = 0.294) or among islands (*Islands*: χ^2^_2_ = 1.22, p = 0.544). We were unable to evaluate differences in abundance of *G. magnirostris* among populations because only two individuals were observed.

The effect of urbanization on the abundance and diversity of the finch community differed among islands (abundance: *Urbanization* × *Island*: χ^2^_1_ = 6.94, p = 0.031; diversity: *Urbanization* × *Island*: χ^2^_1_ = 8.591, p = 0.001; Table S3). Finches were 55% and 18% more abundant in non-urban locations relative to urban populations on Isabela and Santa Cruz, respectively, whereas there was no difference in abundance between urban and non-urban locations on Floreana. We observed a similar trend for diversity (Table S3).

## 4. Discussion

We found that urban environments modify the ecology and evolution of interactions between Darwin’s finches and *T. cistoides*. We measured the effect of urbanization on these interactions using a combination of natural population surveys, field experiments, and finch community observations. Seed and fruit removal rates were higher in urban populations in both natural and experimental populations (Q1), and urbanization modified selection on mericarp morphology (Q2). In natural populations, seed removal imposed stronger selection for large mericarps in urban populations than in non-urban populations, while in experimental populations, fruit removal imposed selection for poorly defended mericarps in urban populations. Lastly, while we found no difference in the abundance of *G. fortis* and *G. magnirostris*, urban environments supported smaller and less diverse ground finch communities than non-urban environments (Q3). Together, our results suggest urbanization can dramatically alter ecological interactions between finches and *T. cistoides*, leading to modified selection on *T. cistoides* populations.

### a) Mericarp predation in urban environments

We observed direct effects of urbanization on seed and mericarp removal. We found that removal was higher in urban populations than in non-urban populations in both the natural and experimental populations. Increased mericarp removal in urban habitats is consistent with the hypothesis that urbanization intensifies interactions between finches and *T. cistoides*. These interactions may have been intensified because *T. cistoides* is more abundant in urban areas on the Galápagos islands (MTJJ and RAJ, personal observation). Humans are the primary dispersers of *T. cistoides* on the Galápagos [27] and *T. cistoides* populations are most likely to be established in and around towns. If ground finches exhibit a functional response to *T. cistoides*, then their consumption of *T. cistoides* seeds may be correlated with the plant’s abundance in a habitat [33]. Such a functional response would explain why urban *T. cistoides* populations experienced greater predation pressure from foraging finches, despite lower finch abundances in these environments.

### b) Selection on mericarp morphology

We found that urbanization alters selection on mericarp morphology. In urban environments, consumption by finches imposed selection for large but poorly defended mericarps. In the natural populations, we observed higher seed removal from small mericarps in urban and non-urban habitats, but this effect was strongest in urban populations. Small mericarps may be more energetically efficient to open [22,34], leading finches to choose small mericarps over large ones. Combined with greater abundances of *T. cistoides* in urban areas, preferential consumption of small mericarps may intensify selection for large mericarps in urban areas. Our results contrast with a previous study which found that finches impose selection for smaller mericarps in natural populations of *T. cistoides* [17]. The differences in findings may be the result of yearly variation in climate that contributes to differences in resource availability. We conducted our study at the end of the dry season following a relatively dry year, but annual variation in precipitation can lead to differences in resource availability [35], and thus differences in the intensity of mericarp consumption by finches. Repeating this study across wet and dry years would help separate out the effects of urbanization from those of climate on the evolutionary ecology of interactions between Darwin’s finches and *T. cistoides*.

In the experimental populations, we observed increased removal of well-defended mericarps from urban populations, whereas in non-urban populations we observed greater removal of mericarps with fewer defences. This result suggests the surprising result that urbanization is associated with finches preferring better defended mericarps. It is presently unclear why urbanization modifies the direction of selection on mericarp defense. Mericarp spines are expected to deter predators from accessing the seeds, an expectation that is consistent with our findings from the non-urban populations and those from Carvajal et al. [17]. However, we found that finches preferred mericarps with longer spines in urban populations. Finches may prefer mericarps with longer spines because they were easier to pick up and manipulate. Alternatively, longer spines can facilitate dispersal by humans [27], but we deliberately placed the dishes in locations where humans were unlikely to disturb them so that mericarp removal would be the result of finch consumption. Further experiments that include finch observations at each experimental site are needed to distinguish between these possibilities.

While our study of natural populations was complementary to our experiments, these two components of our study also differed in several important ways. First, natural populations were available to finches for a longer period of time than the experimental populations, leading to variation in seed removal unrelated to mericarp morphology. Natural populations may have been more likely to experience seed removal because the finches would already have known where to find them. In contrast, finches had to first locate the novel experimental populations before removing the mericarps. Second, because they were exposed longer, the natural populations experienced a greater range of climatic variation, potentially affecting the strength of selection they experienced [35]. Lastly, we were able to track how many seeds were removed from each mericarp in the natural populations, but not from the experimental populations. Seed removal gives a more precise estimate of the fitness effects experienced by the plants and could have resulted in different patterns of selection than mericarp removal.

### c) Urban finch communities

Differences in the urban and non-urban ground finch communities may have contributed to patterns of predation and selection on *T. cistoides*. Finch abundance differed between urban and non-urban sites, although the direction of this difference varied among islands. We observed lower abundance and diversity in urban sites than in non-urban sites on Santa Cruz and Isabela, the largest two islands. This result is consistent with many other studies that find that bird communities are often negatively affected by urbanization (reviewed in [8]). In contrast, there were no significant differences in abundance or diversity between urban and non-urban sites on Floreana, the smallest island, suggesting that town size might play a role in shaping finch communities. Our study tracked how consumption by finches affects the ecology and evolution of *T. cistoides*. To determine if an evolutionary response of *T. cistoides* populations feedbacks to affect the ecology or evolution of finches, studies that combine yearly resampling to track changes in mericarp morphology, beak shape and finch behaviour through time would be necessary.

### d) Conclusion

Together, our results suggest that urbanization modifies the ecology and evolution of interactions between finches and *T. cistoides*. Our study examined the effects of urbanization in an island ecosystem, which are predicted to be particularly sensitive to disturbance. Because of this fragility, perturbations to the landscape through human development and the introduction of invasive species may have large-scale effects on the ecology and evolution of native island species [7]. Our study suggests that urbanization alters the evolutionary ecology of species on islands and identifies trophic interactions as a key mediator of species interactions in the urban island ecosystem.

## Data accessibility

Data and code are included in the supplementary material and will be submitted to the Dryad Digital Repository once accepted for publication.

## Authors’ Contribution

MTJJ and RAJ designed the study and questions, MTJJ, RAJ, and JAC set up the experiment and/or applied for permits, and MTJJ and RAJ collected the data. LRR conducted the analyses and wrote the manuscript, with input from all authors.

## Acknowledgements

We thank the Galapagos National Park for permission and facilitation of our research, and the Charles Darwin Foundation for accommodation and for permitting us to conduct research on their property. C. Richter, M. Johnson, and O. Johnson for help with collecting mericarps and recording data, and D. Reyes and S. Carvajal-Endara for donating data from six natural non-urban populations on Isabela. We thank D. Reyes, and members of the EvoEco Lab for feedback on the manuscript.

## References

1. Grimm NB, Faeth SH, Golubiewski NE, Redman CL, Wu J, Bai X, Briggs JM. 2008 Global change and the ecology of cities. Science 319, 756–760. (doi:10.1126/science.1150195)

2. Seto KC, Guneralp B, Hutyra LR. 2012 Global forecasts of urban expansion to 2030 and direct impacts on biodiversity and carbon pools. Proc. Natl. Acad. Sci. 109, 16083–16088. (doi:10.1073/pnas.1211658109)

3. Niemelä J. 2011 Urban ecology: Patterns, processes, and applications. Oxford: Oxford University Press.

4. Johnson MTJ, Munshi-South J. 2017 Evolution of life in urban environments. Science 358, eaam8327. (doi:10.1126/science.aam8327)

5. Szulkin M, Munshi-South J, Charmantier A. 2020 Urban evolutionary biology. Oxford: Oxford University Press, USA.

6. Rivkin LR et al. 2019 A roadmap for urban evolutionary ecology. Evol. Appl. 12, 384–398. (doi:10.1111/eva.12734)

7. Helmus MR, Mahler DL, Losos JB. 2014 Island biogeography of the Anthropocene. Nature 513, 543–546. (doi:10.1038/nature13739)

8. Aronson MFJ et al. 2016 Hierarchical filters determine community assembly of urban species pools. Ecology 97, 2952–2963. (doi:10.1890/07-1861.1)

9. Vincze E, Seress G, Lagisz M, Nakagawa S, Dingemanse NJ, Sprau P. 2017 Does urbanization affect predation of bird nests? A meta-analysis. Front. Ecol. Evol. 5, 1–12. (doi:10.3389/fevo.2017.00029)

10. Miles LS, Breitbart ST, Wagner HH, Johnson MTJ. 2019 Urbanization shapes the ecology and evolution of plant-arthropod herbivore interactions. Front. Ecol. Evol. 7, 1–14. (doi:10.3389/fevo.2019.00310)

11. Rodewald AD, Kearns LJ, Shustack DP, Applications SE, April N, Kearns J, Shustack P. 2015 Anthropogenic resource subsidies decouple predator-prey relationships. Ecol. Appl. 21, 936–943. (doi:10.1890/10-0863.1)

12. Magle SB, Simoni LS, Lehrer EW, Brown JS. 2014 Urban predator–prey association: coyote and deer distributions in the Chicago metropolitan area. Urban Ecosyst. 17, 875–891. (doi:10.1007/s11252-014-0389-5)

13. Miller CR, Barton BT, Zhu L, Radeloff VC, Oliver KM, Harmon JP, Ives AR. 2017 Combined effects of night warming and light pollution on predator-prey interactions. Proc. R. Soc. B Biol. Sci. 284. 2017119.5 (doi:10.1098/rspb.2017.1195)

14. Grant PR, Grant BR. 2014 40 years of evolution: Darwin’s finches on Daphne Major Island. Princeton, NJ: Princeton University Press.

15. Grant PR. 1999 Ecology and evolution of Darwin’s finches. Princeton, NJ: Princeton University Press.

16. Lack D. 1947 Darwin’s finches. London: Cambridge Univ.

17. Carvajal-Endara S, Hendry AP, Emery NC, Neu CP, Carmona D, Gotanda KM, Davies TJ, Chaves JA, Johnson MTJ. 2020 The ecology and evolution of seed predation by Darwin’s finches on *Tribulus cistoides* on the Galápagos Islands. Ecol. Monogr. 90, 1–17. (doi:10.1002/ecm.1392)

18. Grant PR. 1981 The feeding of Darwin’s finches on *Tribulus cistoides* (L.) seeds. Anim. Behav. 29, 785–793. (doi:10.1016/S0003-3472(81)80012-7)

19. Boag PT, Grant PR. 1984 Darwin’s finches (*Geospiza*) on Isla Daphne Major, Galápagos: Breeding and feeding ecology in a climatically variable environment. Ecol. Monogr. 54, 463–489. (doi:10.2307/1942596)

20. Grant BR, Grant PR. 1982 Niche shifts and competition in Darwin’s finches: G*eospiza conirostris* and congeners. Evolution 36, 637–657. (doi:10.2307/2407879)

21. Boag PT, Grant PR. 1984 The classical case of character release: Darwin’s finches (*Geospiza*) on Isla Daphne Major, Galápagos. Biol. J. Linn. Soc. 22, 243–287. (doi:10.1111/j.1095-8312.1984.tb01679.x)

22. Grant PR, Grant BR. 2006 Evolution of character displacement in Darwin’s finches. Science 313, 224–226. (doi:10.1126/science.1128374)

23. Boag PT, Grant PR. 1981 Intense natural selection in a population of Darwin’s finches (*Geospizinae*) in the Galápagos. Science. 214, 82–85. (doi:10.1126/science.214.4516.82)

24. De León LF, Raeymaekers JAMM, Bermingham E, Podos J, Herrel A, Hendry AP. 2011 Exploring possible human influences on the evolution of Darwin’s finches. Evolution. 65, 2258–2272. (doi:10.1111/j.1558-5646.2011.01297.x)

25. De León LF, Sharpe DMT, Gotanda KM, Raeymaekers JAM, Chaves JA, Hendry AP, Podos J. 2019 Urbanization erodes niche segregation in Darwin’s finches. Evol. Appl. 12, 1329–1343. (doi:10.1111/eva.12721)

26. McNew SM, Beck D, Sadler-Riggleman I, Knutie SA, Koop JAH, Clayton DH, Skinner MK. 2017 Epigenetic variation between urban and rural populations of Darwin’s finches. BMC Evol. Biol. 17, 1–14. (doi:10.1186/s12862-017-1025-9)

27. Johnson MKA, Johnson OPJ, Johnson RA, Johnson MTJ. 2020 The role of spines in anthropogenic seed dispersal on the Galápagos Islands. Ecol. Evol. 10, 1639–1647. (doi:10.1002/ece3.6020)

28. Harvey JA, Chernicky K, Simons SR, Verrett TB, Chaves JA, Knutie SA. 2020 Urban living influences the reproductive success of Darwin’s finches in the Galápagos Islands. bioRxiv. (doi:10.1101/2020.07.08.193623v2)

29. Hendry AP, Grant PR, Grant RB, Ford HA, Brewer MJ, Podos J. 2006 Possible human impacts on adaptive radiation: beak size bimodality in Darwin’s finches. Proc. R. Soc. B Biol. Sci. 273, 1887–1894. (doi:10.1098/rspb.2006.3534)

30. Gotanda KM. 2020 Human influences on antipredator behaviour in Darwin’s finches. J. Anim. Ecol. 89, 614–622. (doi:10.1111/1365-2656.13127)

31. Porter DM. 1971 Notes on the floral glands in *Tribulus* (Zygophyllaceae). Ann. Missouri Bot. Gard. 58, 1–5. (doi:10.2307/2394924)

32. R Development Core Team. 2008 R: A language and environment for statistical computing.

33. Abrams PA. 1982 Functional responses of optimal foragers. Am. Nat. 120, 382–390. (doi:10.1086/283996)

34. Price T. 1987 Diet variation in a population of Darwin’ s finches. Ecology 68, 1015–1028. (doi:10.2307/1938373)

35. Siepielski AM et al. 2017 Precipitation drives global variation in natural selection. Science 962, 959–962. (doi:10.1126/science.aag2773)

36. Instito Nacional de Estadística Y Censos. 2010 Censo De Población Y Vivienda.

