## Supplemental materials for "Urbanization alters ecological and evolutionary interactions between Darwin’s finches and *Tribulus cistoides* on the Galápagos Islands"

**Supplemental methods**

**Study site and system**

The Galápagos Islands are an archipelago 1,000 km off the coast of Ecuador, consisting of hundreds of islands, islets, and small rocks [1]. We studied the effects of urbanization on three of the five inhabited islands in the Galápagos: Floreana, Isabela, and Santa Cruz (Figure 1A). These islands contain towns which differ in area and human population sizes, from 145 people inhabiting 3 km^2^ in Puerto Velasco Ibarra on Floreana, to 2,000 people inhabiting 6 km^2^ in Puerto Villamil in Isabela, and 12,000 people inhabiting 10 km^2^ in Puerto Ayora on Santa Cruz [2], and the population has continued to grow rapidly since the last population census in 2010.

The Galápagos Islands are home to a variety of ground finches (*Geospiza* spp. Gould) which have undergone significant adaptive radiations [3–5]. There are 11 species of ground finches present on the islands [5], including two of interest to our study: the medium ground finch (*G. fortis*) and the large ground finch (*G. magnirostris*). Both of these species are known to feed on *T. cistoides*, although *G. magnirostris* are more adept at extracting seeds [6]. Both species are found on Isabela and Santa Cruz, but only *G. fortis* is present on Floreana because *G. magnirostris* was extirpated over a century ago. *Geospiza fortis and G. magnirostris* are relatively large finches, with large, strong beaks (Figure 1C). Beak morphology is strongly correlated with diet; large beaked finches are capable of consuming larger and harder seeds than small beaked finches [7]. Diet is also associated with temporal variation in seed availability. During wet years, finches feed on seeds from a variety of species, however in dry years, food becomes scarce and large beaked finches feed mostly on hard seeds, such as those produced by *T. cistoides* [8].

*Tribulus cistoides* is thought to be native to tropical and subtropical regions of Africa, but is now widely distributed across tropical continents and islands, including across the Galápagos [9]. On the Galápagos, *T. cistoides* is an herbaceous perennial that grows in arid lowland and coastal habitats [10], where it is often found in sand or volcanic soil alongside roads, trails, and beaches in both urban and non-urban habitats [11]. Most growth of *T. cistoides* occurs during the rainy season, followed by a prolonged flowering period [9]. *Tribulus cistoides* produces hard fruits, which separate into five segments called mericarps. Each mericarp is defended by up to four sharp spines, which serve as a defense against Darwin’s finches [12] and as a mode of seed dispersal [11]. Each mericarp contains 1-7 seeds, which can be accessed by cracking open the mericarp, a feat that is difficult for all but the largest beaked birds [6]. To crack the mericarps, the birds pick it up and apply pressure with their beaks [12]. This process sheers off the ventral side of the mericarp wall, exposing the seeds to removal [12]. This seed removal by finches leaves characteristic damage to mericarps that can be reliably identified even months afterwards.

**Statistical analyses**

*1) Natural population surveys*

We conducted two generalized linear mixed effects models to test for the effects of urbanization on seed removal of *T. cistoides* mericarps in natural populations. Before running the models, we generated composite variables for mericarp size and defense by running a principle component analysis (PCA) on the six morphological variables (Table S1, Figure S1a). We standardized and centered each variable to a mean of 0 and standard deviation of 1 prior to running the PCA. The first principle component (PC) axis was associated with mericarp size and explained 45% of the variation in the data, and all variables loaded in the same direction on this axis (Figure S1a). The second PC axis was associated with defense and explained 20% of the variation in the data. Spine length, position, and presence of lower spine loaded in the same direction on this axis (Figure S1a). Based on these loadings, we used PC1 as a composite representative of size (hereafter: size) and PC2 as a composite representative of defense (hereafter: defense) for the analyses of the natural population survey.

The first model tested for the differences in the number of seeds eaten per mericarp across populations. We ran this model as a Poisson-distributed linear mixed-effects model, using the *glmmTMB* v. 1.0 package [13]. We set the model to account for zero-inflation because more than half the mericarps sampled were uneaten. The model was constructed as follows:

Number of seeds eaten per mericarp ~ *Urbanization*[*U*] + *Island*[*I*] + *Size*[*S*] + *Defense*[*D*] +

*U*:*I* + *U*:*S* + *U*:*D* + *I*:*S* + *I*:*D* + (1|*I*:*Population*)

The number of seeds eaten per mericarp was an integer that ranged from 0-5 and is our proxy for fitness, where more seeds eaten per mericarp results in lower fitness of the plant. *Urbanization* and *Island* were categorical fixed effect variables with two levels (urban and non-urban) and three levels (Floreana, Isabela, Santa Cruz), respectively. *Size* and *Defense* were continuous fixed effect variables that corresponded to the first and second PC axes of mericarp morphology. Mericarps differ in the number of seeds they contain, with bigger mericarps containing more seeds [12]. By including mericarp size (PC1) in the model, we were able to account for differences in seed number due to size, allowing us to more accurately interpret the effects of the other variables in the model. We incorporated population as a random effect in the model and allowed it to vary among islands. This allowed us to account for variation among populations due to differences within islands.

A significant main effect of urbanization or island would indicate that these factors affect the number of seeds eaten per mericarp. A significant main effect of size and/or defense would indicate that these traits influence the number of seeds eaten per mericarp, which would suggest that seed removal imposes selection on these traits. A significant interaction between island and urbanization would indicate that the effect of urbanization on seed removal differs between islands. A significant interaction between urbanization and size/defense would indicate that phenotypic selection on these traits by finches differs between urban and non-urban habitats. Lastly, a significant interaction between island and size/defense would indicate that phenotypic selection on these traits by finches differs between islands.

The second model tested for differences in the proportion of mericarps depredated (whether a mericarp had one or more seeds removed) per population. Although our previous model accounted for variation in the number of seeds produced per mericarp, this second model provided a more conservative estimate of seed removal rate without the potential confounding effect of mericarp size. We ran this model on a binomial distribution as a logistic linear mixed-effects model using the function *glmer* in *lme4* v. 1.1-21 package [14]. The model was constructed similar to the first model:

Proportion of mericarps depredated~ *Urbanization*[*U*] + *Island*[*I*] + *Size*[*S*] + *Defense*[*D*] +

*U*:*I* + *U*:*S* + *U*:*D* + + *I*:*S* + *I*:*D* + (1|*I*:*Population*)

The proportion of mericarps depredated was categorized as a binary variable, where mericarps were recorded as experiencing no seed removal (0) or having at least one seed removed by finches (1). All predictor variables were the same as in the first model, as are the interpretations of effects.

Following model construction, we assessed the assumptions of each generalized linear model by examining the variance inflation factor (VIF) of each variable in the full model using the *vif* function in the *car* v. 3.0-6 package [15]. We found that the VIFs fell below the threshold of 2, suggesting no major collinearity among variables [15]. We assessed the significance of fixed effects from both models using Analysis of Variance (ANOVA) implemented using the *Anova* function in the *car* v. 3.0-6 package [15] to calculate Wald chi-square test statistics with Type III sums-of-squares to test for significant interaction terms. Although Type II sums-of-squares are typically used in cases with incomplete or unbalanced datasets [16], we opted to assess the significance of our results with Type III sums-of-squares because we observed significant interaction terms about which we had *a priori* hypotheses [17,18].

*2) Fruit removal experiment*

We conducted a second logistic regression to determine if urbanization alters selection on mericarp morphology. As with the natural population dataset, prior to analysis we used a PCA to generate composite variables for size and defense (Figure 1D, Table S1, Figure S1). We included the same six morphological measurements as well as mericarp mass. We extracted PC1 (size: 46% of the variation among variables) and PC2 (defense: 17% of the variation among variables) to use in our model; each additional PC axis explained <15% of variation in morphology and were not used in subsequent analyses. These values differ slightly from the PC axes from the natural population analysis because they are extracted from a different set of mericarps. However, PC1 and PC2 have the same direction and magnitude as those from the natural population analysis, thus we are confident that they are capturing similar variation in size and defense.

We tested for differences in mericarp removal from the population using a binomial distributed logistic linear mixed-effects model with a logit link function, implemented using the *glmer* function in *lme4*, as in the previous models. The model was constructed as follows:

Mericarp removal ~ *Urbanization*[*U*] + *Island*[*I*] + *Size*[*S*] + *Defense*[*D*] + *Clipped*[*C*] *+U*:*I* + *U*:*S* + *U*:*D* + *U*:*C* + (1|*I*:*Population*)

Mericarp removal was categorized as a binary variable, where mericarps were recorded either being present in the dish (0) or having been removed from the dish (1). *Urbanization*, *Island*, *Size*, *Defense*, and their interactions were treated and interpreted the same as in the natural population analysis. *Clipped* was a categorical fixed effect factor with two levels (spines removed or intact). A significant main effect term for the clipped treatment would indicate that artificially removing mericarp spines affects the probability of mericarp removal. A significant interaction term between *Urbanization* and *Clipped* would indicate that the effect of spines on susceptibility of fruit removal differs between urban and non-urban environments, and thus urbanization alters selection against mericarp defense. We assessed the assumptions of the model and tested for significance in the same manner as the natural population analysis.

*3)* *Finch community observations*

We assessed variation in the ground finch community using multiple regressions and a constrained ordination analysis. We tested for differences in finch abundance and diversity using multiple linear regression. We constructed the following model:

Abundance/diversity ~ *Urbanization*[*U*] + *Island*[*I*] + *Population* *+ U*:*I*

Abundance was measured as the total number of birds observed at a population. Diversity was calculated as the Shannon Diversity Index, which measures species diversity by combining information on species richness and the evenness in relative abundance of species within a community. *Urbanization*, *Island*, and their interaction were the same as in the previous two analyses. Population was a categorical fixed effect variable used to account for repeated sampling of populations. We determined that the distribution of residuals was normal, and that there was no evidence of multicollinearity among main effects. We assessed the significance of the fixed effects using a Wald chi-square test statistic with Type III sums-of-squares to test for significant interaction terms.

We also measured differences in community composition in response to urbanization using redundancy analysis (RDA). An RDA combines multivariate ordination of a species composition matrix as the response variable with a multiple linear regression of predictor variables, which are fit using a PCA [19]. The result is a series of canonical axes which can be tested for significance using a constrained canonical ANOVA [19].

We conducted the RDA using the *rda* function in the *Vegan* v. 2.5-6 package [20]. The response matrix included species abundance observations with each population listed on a separate line. We transformed the species abundance matrix using a chord transformation to center and standardize the observations [20]. The explanatory matrix included urbanization (urban and non-urban), island, and population. Each line in the explanatory matrix contained data for the population that corresponded to the same line in the response matrix. We tested the significance of the effects using the *anova.cca* function in *Vegan*, with 1000 permutation cycles.

**Supplemental Results**

The results from the proportion of mericarps depredated were largely consistent with those from the number of seeds eaten (Table S4). As with seed removal, the proportion of mericarps attacked declined with mericarp size and was lower on Floreana than either Isabela or Santa Cruz. However, there was no effect of urbanization, defense, or any interactions on the proportion of mericarps depredation.

The results from the RDA were consistent with the results for abundance and diversity (Figure S2; Table S3). The composition of the finch community differed between urban and non-urban habitats (*Urbanization*: F_1,51_ = 8.49, p < 0.001) and among islands (*Island*: F_2,51_ = 4.18, p < 0.001), as did the effect of urbanization among islands (*Urbanization* × *Island*: F_2,51_ = 2.71, p = 0.023). Although communities somewhat overlapped across populations, small ground finches (*G. fuliginosa*) were more frequently observed in urban populations than in non-urban populations.

**Literature Cited**

1.	Geist D. 1996 On the emergence and submergence of the Galapagos Islands. *Not. Galapagos* , 5–9.

2.	Censos INDEY. 2010 Censo De Población Y Vivienda.

3.	Grant BR, Grant PR. 1982 Niche Shifts and Competition in Darwin’s Finches: Geospiza conirostris and Congeners. *Evolution (N. Y).* **36**, 637–657. (doi:10.2307/2407879)

4.	Grant PR, Grant BR. 2006 Evolution of character displacement in Darwin’s finches. *Science (80-. ).* **313**, 224–226. (doi:10.1126/science.1128374)

5.	Lamichhaney S *et al.* 2015 Evolution of Darwin’s finches and their beaks revealed by genome sequencing. *Nature* **518**, 371–375. (doi:10.1038/nature14181)

6.	Grant PR. 1981 The feeding of darwin’s finches on *Tribulus cistoides* (L.) seeds. *Anim. Behav.* **29**, 785–793. (doi:10.1016/S0003-3472(81)80012-7)

7.	Abbott I, Abbott LK, Grant PR. 1977 Comparative Ecology of Galapagos Ground Finches (Geospiza Gould): Evaluation of the Importance of Floristic Diversity and Interspecific Competition. *Ecol. Monogr.* **47**, 151–184. (doi:10.2307/1942615)

8.	Grant PR, Grant BR. 1980 Annual variation in Finch numbers, foraging and food supply on Isla Daphne Major, Galápagos. *Oecologia* **46**, 55–62. (doi:10.1007/BF00346966)

9.	Porter DM. 1971 Notes on the floral glands in Tribulus Zygophyllaceae. *Ann. Missouri Bot. Gard.* **58**, 1–5.

10.	Schweickerdt HG. 1868 An account of South African species of *Tribulus Tourn. ex Linn* . **36**.

11.	Johnson MKA, Johnson OPJ, Johnson RA, Johnson MTJ. 2020 The role of spines in anthropogenic seed dispersal on the Galápagos Islands. *Ecol. Evol.* **10**, 1639–1647. (doi:10.1002/ece3.6020)

12.	Carvajal-Endara S, Hendry AP, Emery NC, Neu CP, Carmona D, Gotanda KM, Davies TJ, Chaves JA, Johnson MTJ. 2020 The ecology and evolution of seed predation by Darwin’s finches on *Tribulus cistoides* on the Galápagos Islands. *Ecol. Monogr.* **90**, 1–17. (doi:10.1002/ecm.1392)

13.	Brooks ME, Kristensen K, van Benthem KJ, Magnusson A, Berg CW, Nielsen A, Skaug HJ, Mächler M, Bolker BM. 2017 glmmTMB balances speed and flexibility among packages for zero-inflated generalized linear mixed modeling. *R J.* **9**, 378–400. (doi:10.32614/rj-2017-066)

14.	Bates D, Maechler M, Bolker B, Walker S. 2015 Fitting Linear Mixed-Effects Models Using {lme4}. *J. Stat. Softw.* **67**, 1–48.

15.	Fox J, Weisburg S. 2011 *An R Companion to Applied Regression*. Second Ed. Thousand Oaks CA: Sage.

16.	Langsrud Ø, June R, As TIII, Iii T, Type T, Ii T, Iii TT, Ii TT. 2003 ANOVA for unbalanced data: Use Type II instead of Type III sums of squares. *Stat. Comput.* , 163–167.

17.	Shaw RG, Mitchell-olds T. 1993 Anova for unbalanced data: An overview. *Ecology* **74**, 1638–1645.

18.	Hector A, von Felten S, Schmid B. 2010 Analysis of variance with unbalanced data: An update for ecology & evolution. *J. Anim. Ecol.* **79**, 308–316. (doi:10.1111/j.1365-2656.2009.01634.x)

19.	Legendre P, Legendre L. 2012 Canonical analysis. In *Developments in Environmental Modelling*, pp. 625–710. Elsevier.

20.	Oksanen J, Kindt R, Legendre P, O’Hara B, Simpson GL, Solymos PM, Stevens MHH, & Wagner H. 2008 The vegan package. *Community Ecol. Packag.* , 190.

**Supplemental Tables**

Table S1: Proportion of variance explained by each PC axis for size and defense traits of mericarps from the natural populations and experimental populations.

|  | Proportion of Variance | |
| --- | --- | --- |
|  | Natural populations | Experimental populations |
| PC1 | 0.4472 | 0.4558 |
| PC2 | 0.2005 | 0.1700 |
| PC3 | 0.1660 | 0.1416 |
| PC4 | 0.0899 | 0.1213 |
| PC5 | 0.0647 | 0.0668 |
| PC6 | 0.0317 | 0.0297 |

Table S2: Results from the natural population survey and mericarp removal experiment across three islands in the Galápagos. We examined two response variables: the number of seeds eaten per mericarp (natural population results; *N* = 1699 mericarps) and the proportion of mericarps which were removed from a site (experiment results; *N* = 2120 mericarps). Results presented in the table show the fixed effects of urbanization, island, mericarp size and defense, spine clipping (experiment results only), and their interactions. The significance of fixed effects was estimated with Wald χ^2^ test statistics using Type III sums-of-squares.

|  |  | A) Seeds eaten per site | | B) Mericarp removal | |
| --- | --- | --- | --- | --- | --- |
|  | df | χ^2^ | P-value | χ^2^ | P-value |
| Urbanization | 1 | 3.91 | **0.048** | 4.98 | **0.026** |
| Island | 2 | 3.98 | 0.137 | 2.91 | 0.234 |
| Size | 1 | 10.74 | **< 0.001** | 2.40 | 0.121 |
| Defense | 1 | 3.13 | 0.077 | 0.46 | 0.495 |
| Clipped | 1 | NA | NA | 8.44 | **0.004** |
| Urbanization × Island | 2 | 5.77 | 0.056 | 3.28 | 0.194 |
| Urbanization × Size | 1 | 4.51 | **0.034** | 2.25 | 0.134 |
| Urbanization × Defense | 1 | 2.51 | 0.113 | 4.24 | **0.039** |
| Urbanization × Clipped | 1 | NA | NA | 0.05 | 0.823 |
| Island × Size | 2 | 8.89 | **0.012** | 5.74 | 0.057 |
| Island × Defense | 2 | 6.69 | **0.035** | 2.46 | 0.292 |

Table S3: Results from finch surveys across islands (*N* = 60 observations) for abundance, Shannon diversity, and community composition. Results presented in the table show the fixed effects of urbanization, island, population, and the interaction between urbanization and island. The significance of fixed effects for abundance and diversity were estimated using Wald χ^2^ test statistics with Type III sums-of-squares. The significance of fixed effects for community composition were estimates with pseudo-F test statistics.

| Response | Predictor | df | χ^2^/F-value | P-value |
| --- | --- | --- | --- | --- |
| Abundance | Urbanization | 1 | 1.73 | 0.189 |
|  | Island | 2 | 23.10 | **< 0.001** |
|  | Population | 5 | 15.84 | **0.007** |
|  | Urbanization × Island | 2 | 6.94 | **0.031** |
| Diversity | Urbanization | 1 | 3.00 | 0.090 |
|  | Island | 2 | 2.10 | 0.134 |
|  | Population | 5 | 1.47 | 0.218 |
|  | Urbanization × Island | 2 | 8.59 | **0.001** |
| Community | Urbanization | 1 | 8.49 | **< 0.001** |
| Composition | Island | 2 | 4.18 | **0.002** |
| (RDA) | Population | 5 | 0.41 | 0.971 |
|  | Urbanization × Island | 2 | 2.71 | **0.023** |

Table S4: Results from the natural population surveys across three islands in the Galápagos with the response as the proportion of mericarps which were subjected to seed removal (*N* = 1699 mericarps). Results presented in the table show the fixed effects of urbanization, island, mericarp size and defense, and their interactions. The significance of fixed effects were estimated with Wald χ^2^ test statistics using Type III sums-of-squares.

|  | DF | χ^2^ | P-value |
| --- | --- | --- | --- |
| Urbanization | 1 | 1.80 | 0.180 |
| Island | 2 | 7.71 | **0.021** |
| Size | 1 | 2.20 | 0.138 |
| Defense | 1 | 2.25 | 0.134 |
| Urbanization × Island | 2 | 2.42 | 0.298 |
| Urbanization × Size | 1 | 1.56 | 0.212 |
| Urbanization × Defense | 1 | 0.84 | 0.359 |
| Island × Size | 2 | 5.15 | 0.076 |
| Island × Defense | 2 | 12.12 | **0.002** |

Figure S1: Loadings of size and defense traits of mericarps from (a) natural and (b) experimental populations (b). Each variable is colour-coded by their contribution to the loadings.

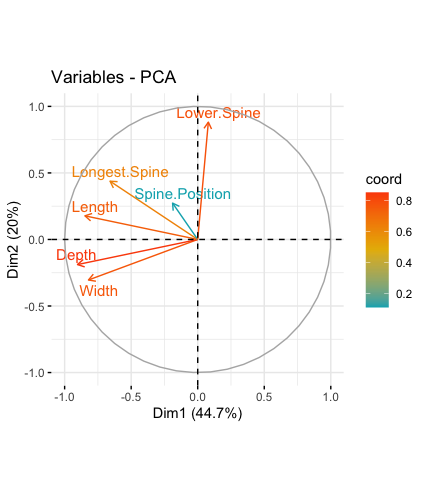

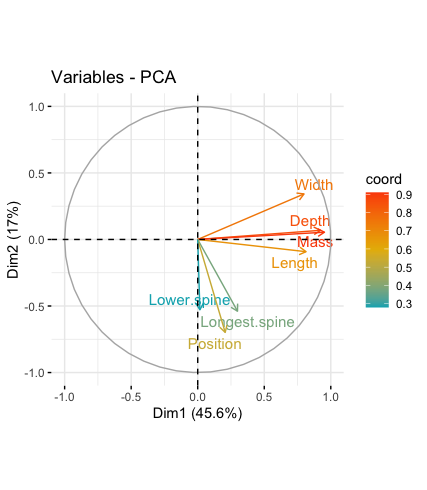

a

b

Figure S2: Community composition plot of urban (red) and nonurban (black) finches across the three islands in our study.

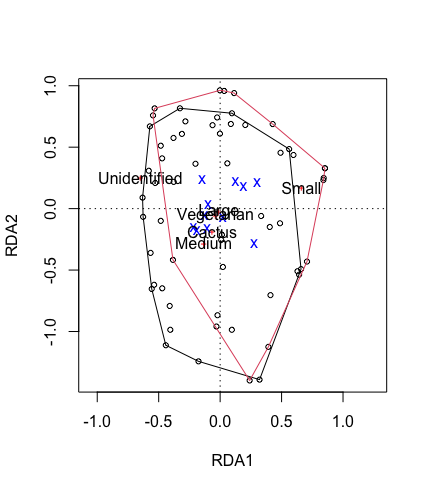
